# Polyelectrolyte brush bilayer under shear at linear and nonlinear response regimes: A combination of the density functional theory framework and the scaling theory

**DOI:** 10.1101/2021.05.19.444845

**Authors:** Mike J. Edwards

## Abstract

The density functional theory framework and the scaling theory are employed to approach the problem of the Polyelectrolyte brush bilayer under shear. It turns out that, the system at shear rates larger than a critical shear rate undergo a global restructuring during which chains stretch in the shear direction. In the absence of the electrostatic interactions as well as the hydrodynamic interactions, this global restructuring causes a sublinear scaling of the shear stress with the shear rate which makes the shear thinning effect. Nevertheless, in the presence of the hydrodynamic interactions, not only there is no sublinear regime but also a weak superlinear regime which makes a weak shear thickening effect. In the presence of the electrostatic interactions, the stress tensor components change by their second Virial coefficients, however, their shear rate power law are untouched. Nonetheless, the kinetic friction coefficient is independent of the electrostatic interactions. This suggests that the lubrication is not very much different than the neutral bilayers and the electrostatic interactions do not change that. The results of this study offers that maybe nature uses another mechanism to reduce friction coefficient in synovial joint and other biological systems.

**SIGNIFICANCE:** The significance of this study is that it strongly criticizes the theoretical approach to the same system which is already published (T. Kreer, *Soft Matter*, **12**, 3479 (2016)). Moreover, the results of this study may help our understanding from Biological systems and optimization of artificial synovial joint which is the core of this study.

## INTRODUCTION

The building blocks of life are linear macromolecular structures which are polymers (1). These macromolecular structures are made by connecting around 10^4^ to 10^5^ repeating units or monomers to each other. Monomers are connected to each other by covalent bonds, in which, the atoms share their valence electrons. Polymers could be found in nature in a varieties of structures by connecting linear segments to each other. From polymer brushes which are made by grafting linear chains onto a surface all the way to polymer stars that are made by fixing linear chains in one point (2–5). In nature, there are more sophisticated polymer structures like dendrimers that are polymer stars with branching arm chains and gels which are a randomly connected bulk of chains (2–5).

Polymer brushes form when a large number of linear chains are densely grafted to (from) a surface (2–5). The steric repulsion between the nearby chains, stretches the chains in the perpendicular direction. One of the most important features of the polymer brushes is in lubrication (2–5). It has been known for decades, that polymer brush covered surfaces exhibit fascinating lubrication properties. This super lubricity comes from the fundamental molecular structures of this system under shear motion. Some researchers believe that the interpenetration between the brushes is responsible for this super lubricity (6). They relate the sublinear regime which appears in the shear thinning effect to the vanishing of the interpenetration zone between brushes (6). The most obvious criticize to this idea would be to ask why the shear stress does not vanish as the interpenetration zone does. The most straight answer to this question would be because vanishing of the interpenetration zone does not cause the shear thinning effect. In contrast, the chains which get stretched strongly in the shear direction and the system undergoes a global restructuring is the key effect (7). In the present study, I shall address these issues by extrapolating my theory to the Polyelectrolyte brush bilayers (PEBs) in the presence of the hydrodynamic interactions (HIs).

## METHOD

Polymer brush bilayers (PBBs) are shown schematically in Fig. (1). Linear polymer chains with the degree of polymerization *N*, and the *Kuhn* length *a* are grafted to two parallel opposing surfaces. The grafting density is *σ* and distance between the surfaces is *D*. To obtain the equilibrium properties of the PBBs, one could use the DFT framework that has been described in detail in (8). The equation of state of the PBBs as well as brush extensions are the quantities of interest.

**Figure 1:**
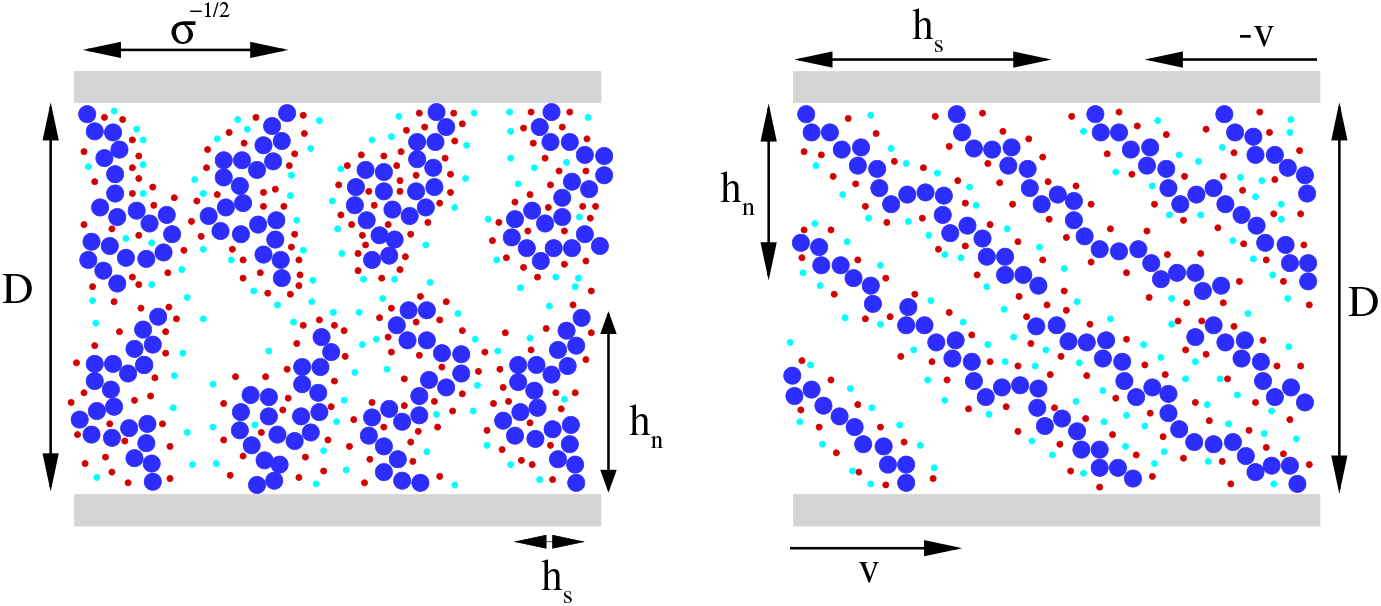
Two dimensional sketches of the PBB system with CIs. Left: The PBB at equilibrium conditions. Blue particles are monomers, the red particles are oppositely charged CIs and the light blue particles are likely charged particles. Distance between surfaces is *D*, the average distance between grafted chains is *σ*^−1/2^, average lateral height of chains is *h_s_* and the average perpendicular height of chains is *h_n_*. Right: The PBB system at nonstationary shear conditions. The shear velocity of surfaces is *v*.

Assume that a fluid with viscosity *η* is confined between two parallel flat surfaces with distance *D*. The systems under shear are categorized as a system under special flow, that is shear. The governing differential equation of systems under flow at small scales is the Stokes equation (9). The Stokes equation for shear flow reduces to an ordinary differential equation and the elements of the stress tensor become 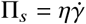 and Π_*n*_ = *p*. Where, 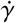 is the shear rate and *p* the equation of state. The equation of state is obtained through DFT, so, one needs to calculate the viscosity to have the complete stress tensor of the PBBs.

The viscosity of a system is an intrinsic property of that system and it has roots in fundamentals at the molecular scale. Essentially the viscosity is defined as the multiplication of equation of state *p* and the collision time *τ*. The collision time is the time during which a particle travels between two subsequent collisions. In the other words, the collision time is the duration of a mean free path. The collision time for the PBBs is calculated as mean free path squared divided by diffusion constant (7, 10–12).

The hydrodynamic interactions (HIs) between particles in a system loom at very low frequencies which is called *hydrodynamic limit* (10). In the hydrodynamic limit which could be seen in dilute and semidilute systems, the particles interact through *Oseen tensor* elements (11). By taking into account the HIs, the particles diffusion constant is modified. This modification takes place in the way that, the diffusion constant is now a function of the solvent viscosity and not the monomer friction constant (11). Therefore, by using a different diffusion constant, one obtains a different collision time and a different viscosity.

In a Polyelectrolyte solution, the monomers interact through electrostatic interactions (EIs). However, once a Polyelectrolyte is suspended in a solvent like water (which is the universal solvent), the oppositely and likely charged free particles, counterions (CI), are dissociated into the solvent as well. The presence of CIs in the solvent causes the screening of the EIs everywhere in the system. So, the screened EIs are less long range than bare EIs. The theory of screened EIs are developed by Debye and Hückel (13). The presence of the EIs in a system is presented in the second Virial coefficient and they do not influence the diffusion constant and the collision time.

## RESULTS

### The PBB under shear in the absence of the hydrodynamic interactions

The PBBs under shear at small shear rates respond linearly to the shear forces. It means that the shear forces can not ovecome thermal forces and the equation of state does not change. In this linear response regime, the system is governed by the *Rouse dynamics* (11). In the Rouse dynamics, the shear rate below 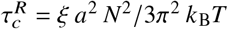 belong to the linear response regime. In the shear rates above this critical time scale, the chains start stretching in the shear direction and the nonstationary shear regime looms. To present the critical shear rate, the dimensionless quantity, Weissenberg number 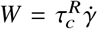 is introduced. So, at *W* ≫ 1 the system resides in the linear response regime and at *W* ≪ 1 it resides in the nonlinear response regime. The DFT framework calculations show that, at linear regime, the shear and normal chain extensions are given as follows (7, 8),

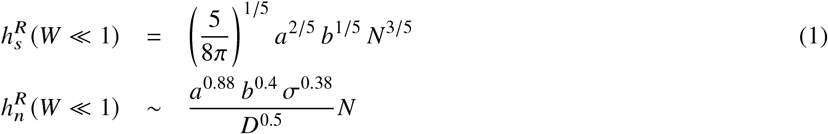

The DFT framework calculations, also, show that the shear and normal stress components at linear response regime are given as follows (7). Note that, to calculate the shear stress one need to calculate the intrinsic viscosity which needs the collision time. The collision time in the Rouse dynamic is given as (*D*/2*σ*)^2/3^ (*ξN*^1/3^/*k*_B_*T*) (7, 10, 11).

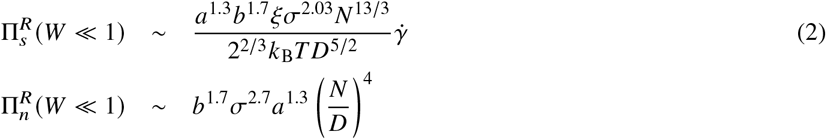

After calculating the physical quantities at linear response regime, one can obtain the nonlinear response regime quantities by using phenomenological argument and constructing an scaling function. The scaling function is considered to be in the form of Ξ *W* = *W^α^* and it multiplied by the physical quantities at linear response regime. The phenomenological argument helps to determine the exponent *α*. In the case of the PBBs under shear, the phenomenological argument suggests that the chains stretch in the shear direction at high shear rates. This means that, the universal power law of the physical quantities in terms of the degree of polymerization between shear and normal components are swapped. In the case of the chain extensions, the shear chain extension must be scaled as *N* at high shear rates which leads to *α* = 4/5 and the normal chain extension must be scaled as *N*^3/5^ at high shear rates which leads to *α* = −1/5. The simplified chain extensions at high shear rates are given as follows,

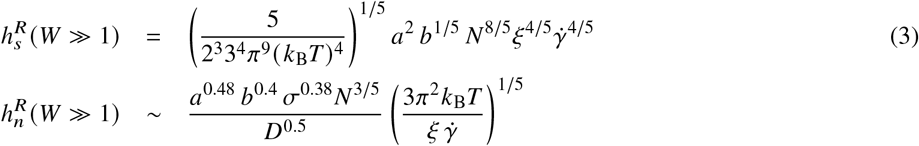

In the case of the stress tensor components, the phenomenological argument is that the shear stress must be scaled as *N*^4^ at high shear rates leading to *α* = −1/6. Likewise, the normal stress must be scaled as *N*^13/3^ which leads to *α* = 1/6. The stress tensor components at nonlinear response regime are given as follows,

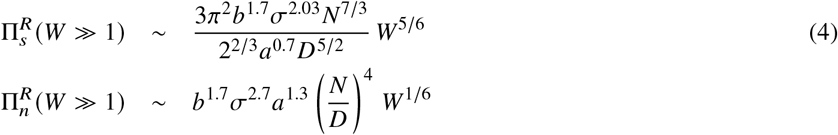

The kinetic friction coefficient could be calculated through dividing the shear stress by the normal stress as follows,

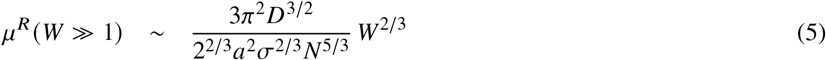

The viscosity at high shear rates could be calculated through dividing the shear stress at high shear rates by *W* which is given as follows,

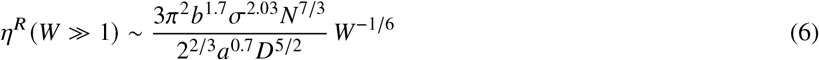

### The PBB under shear in the presence of the hydrodynamic interactions

When the density of the system is close to dilute or semidilute, the particles feel the low frequency forces which are in the hydrodynamic limit (10). In the hydrodynamic limit, the particles interact via HIs. The effect of HIs are represented in the diffusion constant of the particles (11). In the systems with HIs, the *Zimm dynamics* governs the dynamics of the system (11). In this conditions, the collision time of the system is given as (*D*/2*σ*)^2/3^ (*η_s_a*/*k*_B_*TN*^1/15^). The phenomenological argument to construct the scaling function is that the shear chain extension must be scaled as *N* leading to *α* = 2 9 and the normal chain extension must be scaled as *N*^3/5^ which leads to *α* = −2/9 at high shear rates. The simplified chain extensions at nonlinear response regime with HIs is given as follows. Note that, the chain extensions at linear response regime are equal to the those in the system in the absence of HIs.

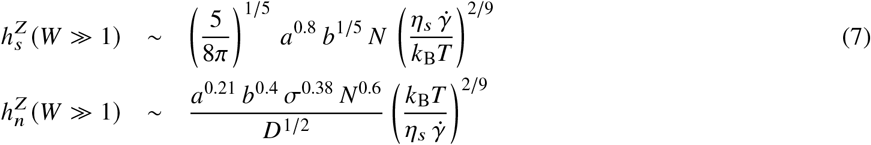

The components of the stress tensor at low shear rates, could be calculated as follows. Note that, the shear stress is influenced by the HIs due to diffusion constant, however, the normal stress is not changed.

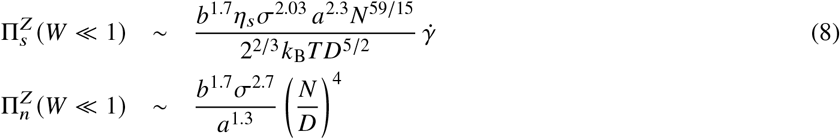

The phenomenological argument to calculate the shear stress at high shear rates is that it must be scaled as *N*^4^ leading to *α* = 1/27. Likewise, the normal stress at high shear rates must be scaled as *N*^59/15^ leading to *α* = −1/27. The simplified stress tensor components for nonlinear response regime are given as follows,

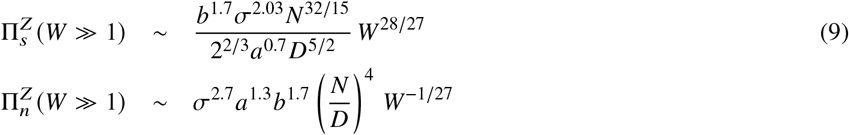

As it is seen, the shear stress is scaled as 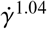. This shows that the PBB system in the presence of the HIs, does not follow a sublinear regime and it is weakly tends toward the superlinear regime. Dividing the shear stress by the normal stress one could get the following expression for the kinetic friction coefficient at high shear rates,

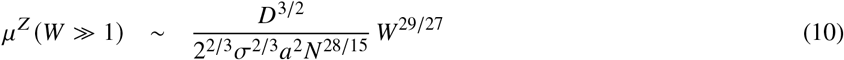

To obtain the viscosity at high shear rates, one could again divide the shear stress at high shear rates by the Weissenberg number,

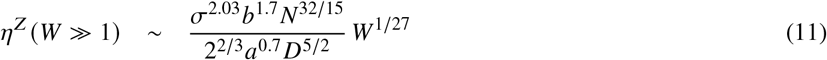

### The PEB under shear in the presence of the hydrodynamic interactions

Particles in the PEBs in the presence of the HIs experience both EIs and HIs. This is very close to the realistic situations which occur in experiments. The EIs modify the second Virial coefficient and the HIs make major modifications on the system parameters. In Fig. (2), the second Virial coefficient is shown in terms of the Bjerrum length. It is seen that the second Virial coefficient is larger in the presence of the EIs and increasing the Bjerrum length makes it closer to the neutral systems. Therefore, the EIs does not alter the universal power law of the shear rate. It means that the EIs has nothing to do with quality of the shear thinning or thickening effects. Consequently, the results of the HIs section hold in this section as well with difference their second Virial coefficient. Since, the kinetic friction coefficient is independent of the second Virial coefficient, therefore, theoretically, the EIs does not alter the friction between the surfaces.

**Figure 2:**
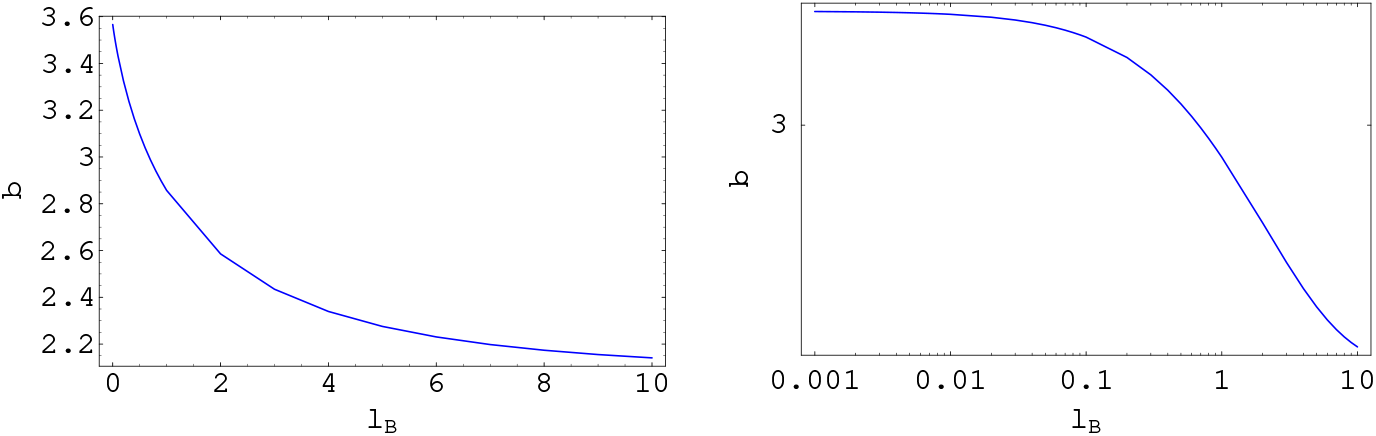
The second Virial coefficient in terms of the Bjerrum length. Left: The linear plot. Right: The logarithmic plot.

## CONCLUDING REMARKS

In this study, the problem of a PEB under stationary shear motion is presented. The results are shown for both linear and nonlinear regimes. Also, the results are promising by exhibiting a good combination of the DFT framework and the scaling theory. The shear thinning effect is observed in the system without HIs, however, there is no sign of this effect with HIs. Moreover, the HIs drive the system towards a weak shear thickening effect. This phenomenon apparently is due to the fact that HIs are repulsive and make system to look more melt like. One of the most remarkable results of this study is that it shows the EIs has no role in the lubrication between surfaces. Therefore, it suggests that maybe the nature, uses another mechanism to decrease the friction in the synovial joints. Another important issue here is that, the blob picture that has been in the work already published about PEBs, apparently does not predict the correct aspects of this system and a major review must be done on that. Mainly, in the present study there is no sign of correlation between lubrication and the EIs. Nonetheless, the blob picture predicts a melt like behavior for PEBs.

## ACKNOWLEDGMENTS

I thank the Cold Spring Harbor Laboratory (CSHL) and the Regeneron Pharmaceuticals Inc. for financial support, Wolfram Research for Mathematica, the Red hat for the Fedora Linux, the Tecplot Inc. for Tecplot visualization software, the Weizmann Institute of Science for XMGrace, and the Overleaf latex editor.

## REFERENCES

1. Rubinstein, M., and R. Colby, 2004. Polymer Physics. OUP Oxford, Oxford, first edition.

2. Advincula, R., W. Brittain, K. Caster, and J. Rühe, 2006. Polymer Brushes: Synthesis, Characterization and Applications. Wiley. https://books.google.com/books?id=hMtdlxLyQJ4C.

3. Azzaroni, O., and I. Szleifer, 2017. Polymer and Biopolymer Brushes: for Materials Science and Biotechnology. Wiley. https://books.google.com/books?id=W8Q_DwAAQBAJ.

4. Lilge, I., 2017. Polymer Brush Films with Varied Grafting and Cross-Linking Density via SI-ATRP: Analysis of the Mechanical Properties by AFM. BestMasters. Springer Fachmedien Wiesbaden. https://books.google.com/books?id=UPs1DwAAQBAJ.

5. Mittal, V., 2012. Polymer Brushes: Substrates, Technologies, and Properties. CRC Press. https://books.google.com/books?id=9EfNBQAAQBAJ.

6. Kreer, T., 2016. Polymer-brush lubrication: a review of recent theoretical advances. Soft Matter 12.

7. Edwards, M., 2019. Polymer brush bilayers under stationary shear motion at linear response regime: A theoretical approach. bioRxiv https://www.biorxiv.org/content/early/2019/03/04/565796.

8. Edwards, M., 2020. Polymer brush bilayers at thermal equilibrium: A theoretical approach. bioRxiv https://www.biorxiv.org/content/early/2020/11/22/316141.

9. Safran, S., 2003. Statistical Thermodynamics Of Surfaces, Interfaces, And Membranes. Boca Raton: CRC Press., Oxford, first edition.

10. Schwabl, F., and W. Brewer, 2006. Statistical Mechanics, Advanced Texts in Physics. Springer Berlin Heidelberg, Oxford, first edition.

11. Doi, M., and S. F. Edwards, 1986. Theory of polymer dynamics. Oxford science publication, Oxford, first edition.

12. Gennes, P. G., 1979. Scaling Concepts in Polymer Physics. Cornell University Press, Oxford, first edition.

13. Debye, P., and E. Hüeckel, 1923. Zur Theorie der Elektrolyte. I. Gefrierpunktserniedrigung und verwandte Erscheinungen. Physikalische Zeitschrift. 24.

